# Effect of Mangiferin Isolated from *Hedysarum neglectum* L. Root Culture on Fat Accumulation in Experiments *In Vivo*

**DOI:** 10.1101/2023.05.10.540104

**Authors:** Anastasia M. Fedorova, Alexander Y. Prosekov, Irina S. Milentyeva, Varvara I. Minina, Elena V. Ostapova

## Abstract

Plant polyphenols possess diverse medicinal and therapeutic properties. For instance, they can prevent such age-related diseases as stroke, heart attack, type II diabetes, and their various pathological complications. Mangiferin is a polyphenolic compound with a C-glycosylxanthone structure. This biologically active substance is known for its antioxidant, anti-inflammatory, antimicrobial, antidiabetic, and immunomodulatory properties. This research features mangiferin isolated from *Hedysarum neglectum* L. root culture and its effect on fat accumulation in *Caenorhabditis elegans* N2 Bristol. A set of experiments *in vivo* proved that mangiferin was able to inhibit fat accumulation in the model organism. The optimal concentration was determined as 50 μm: it provided 44.5% fluorescence.

## Introduction

Aging is a complex process associated with a gradual decline in cellular and physiological functions. The recent achievements of healthcare and medical sciences have prolonged human life expectancy, and the proportion of people over 65 continues to increase [1]. Unfortunately, chronic age-related diseases, such as stroke, heart attack, and type II diabetes, remain quite common. Therefore, understanding the key mechanisms of aging and age-related diseases is an important issue. Obesity is a complex multifactorial problem. Body weight depends on the environment, genetic factors, and energy imbalance. However, socioeconomic conditions can also trigger obesity [2]. Aging is associated with an increase in abdominal white adipose tissue and deposition of fat in skeletal muscles, which affects insulin sensitivity [3]. After retirement, people usually experience significant changes in their lifestyle, which often leads to a chronic positive energy balance, obesity, and, eventually, age-related diseases.

Model organisms, e.g., *Caenorhabditis elegans*, make it possible to study such processes as obesity in laboratory conditions. *C. elegans* is a nematode worm with a relatively short life span that can produce up to 300 offspring and has a generation time of three days, which makes it a perfect model organism for experiments *in vivo* [4, 5]. *C. elegans* are much cheaper and easier to maintain than many other model organisms because the worm is approximately 1 mm in length. *C. elegans* feeds on such affordable foodstuffs as *Escherichia coli* and can be stored for a long time at -80 °C or in liquid nitrogen [6]. *C. elegans* can undergo complex genetic modifications for rapid, large-scale, and high-throughput screening to test new medicines or individual bioactive compounds [7]. This model organism is also suitable for antioxidant, anti-aging, and neuroprotective studies.

Polyphenols possess hepatoprotective, antioxidant, hypoglycemic, and other beneficial properties [8]. They improve the metabolic activity of anti-obesity foods that prevent or inhibit obesity [9]. Mayneris-Perxachs *et al*. [10] added hesperidin and nargenin to cookies, which improved the metabolic syndrome in obese rats and reduced their body weight, total body fat, total cholesterol, and oxidative stress.

Mangiferin (xanthone) is a polyphenolic compound that can be extracted from *Hedysarum neglectum* L. [11]. Mangiferin is known for its antiviral and antioxidant properties [12]. Extracts with mangiferin proved to be quite potent antioxidants (EC_50_: 5.8 ±0.96 μg/mL). They also demonstrated hepatoprotective activity against carbon-tetrachloride-induced liver damage: their radical purification system proved quite effective *in vivo* [13]. Mangiferin had a dose-dependent ability to reduce hydrogen-peroxide-induced oxidation and lipid oxidation in human peripheral blood lymphocytes [14]. Its beneficial properties make mangiferin an excellent anti-obesity drug. The research objective was to study the effect of mangiferin isolated from *Hedysarum neglectum* L. root cultures on fat accumulation in *C. elegans*.

## Objects and methods

The research featured mangiferin isolated *in vitro* from the extract of *Hedysarum neglectum* L. root culture.

The root culture was obtained at earlier stages as described by Dyshlyuk *et al*. [11]. The roots were treated with *Agrobacterium rhizogenes* 15834 Swiss (Moscow, Russia). After 12-48 h of exposure to the bacterial suspension, the explants were rinsed in sterile water, dried with sterile filter paper, and placed on Gamborg B5 solid medium that contained 500 mg/L Claforan (Patheon, UK). The explants were cultivated on the solid medium supplemented with 250 mg/l Claforan and without hormones. The root culture was cultivated *in vitro* in the dark at +23 °C on an LSI-3016A shaking incubator at 100 rpm for five weeks.

The root culture extract of *Hedysarum neglectum* L. corresponded with the extraction parameters defined by Faskhutdinova *et al*. [15]: 6 h, 30°C, and 70% ethanol.

The procedure of isolating mangiferin from the extract was described in [15] and followed the scheme in Figure 1. The mangiferin purification degree was ≥95%.

**Figure 1.**
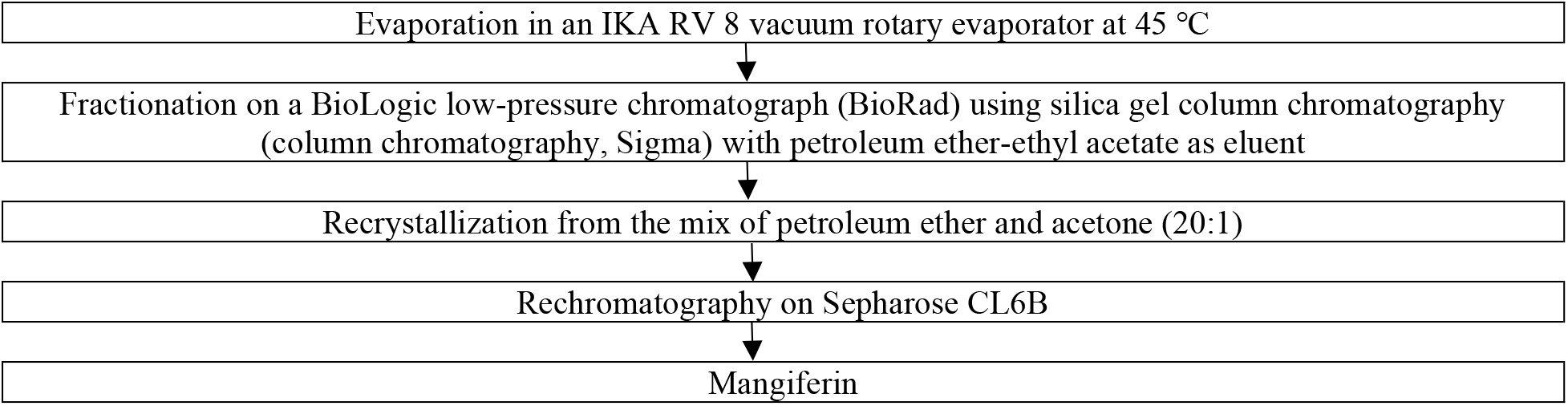
Isolating margiferin from *Hedysarum neglectum* L. root culture ethanol extract

A wild strain of nematode *Caenorhabditis elegans* N2 Bristol served as a model organism to study the effect of mangiferin on fat accumulation. The samples were provided by the Laboratory for Innovative Drugs and Agrobiotechnologies, Moscow Institute of Physics and Technology, National Research University (Dolgoprudny, Russia).

*C*.*elegans* fed on *Escherichia coli* OP50 provided by the Engelhardt Institute of Molecular Biology, Russian Academy of Sciences (Moscow, Russia). The bacteria were added once at the onset of the experiment to a final concentration of 5 mg/mL.

The nematodes were cultivated on solid nematode growth medium (NGM) agar which included 1000 ml ddH_2_O, 2.5 g bactopeptone, 3 g NaCl, and 17 g bactoagar. It was autoclaved at 120 °C for15 min and mixed with 1 ml 1M CaCl_2_, 1 ml 1M MgSO_4_, 1 ml 5 mg/ml cholesterol, and 25 ml K_2_HPO_4_+ KH_2_PO_4_ at pH 6.0. First, we put 50 μl of *E. coli* OP50 overnight culture into the *center* of a 100 mm sterile Petri dish. Then, we used a sterile glass rod to draw a square in the center of the dish with a drop of bacterial culture, without touching the walls. After that, we incubated the plates for 24 at 37 °C in a climate chamber (Binder, Germany). Finally, we transferred the nematodes onto the NGM agar plates. A transfer loop was glowed in fire and cooled in the agar to hook a nematode and transfer it under sterile conditions to the bacterial lawn in the center of a new NGM agar plate. A 0.5 cm × 0.5 cm piece of agar with nematodes was cut out of the NGM dish, transferred to a new dish, and placed top surface down in the center to be incubated at 20°C.

To synchronize the nematodes, we added 5-10 ml of sterile water to the agar Petri dish with the nematodes and pipetted it several times to free all nematodes and their eggs from the agar. The liquid was transferred from the Petri dish to a 50 ml centrifuge tube and centrifuged at 1200 rpm (280 g) for 2 min. After that, we removed the supernatant, washed the precipitate with 10 ml of distilled water, and repeated the procedure. Then, we mixed 1 ml 10 n. NaOH with 2.5 ml sodium hypochlorite and 6.5 ml water and added the mix to the sediment. The hydrolysis of nematodes underwent a microscopy while being stirred and vortexed for 5 min with a break every 2 min. To neutralize the reaction, we prepared 5 ml of M9 medium, which included 1000 ml ddH_2_O, 3 g KH_2_PO_4_, 5 g NaCl, 6 g Na_2_HPO_4_, and 1 ml of 1 M MgSO_4_. The mix was centrifuged at 2500 rpm (1100 g) for 2 min. After removing the supernatant, we resuspended the pellet in 10 ml of fresh sterile water and triplicated the neutralization and centrifugation. The precipitate was washed four times with 10 ml of S medium, which included 1000 ml S base, 10 ml 1 M potassium citrate pH 6, 10 ml trace metal solution, 3 ml 1 M CaCl_2_, and 3 ml 1 M MgSO_4_. Finally, we removed the supernatant, added 10 ml of S medium, and left the test tube with nematode eggs on a shaker at room temperature for 24 h until the nematodes entered L1 stage.

To cultivate the nematodes in a liquid medium, we added the overnight bacterial culture of *E. coli* OP50 to the L1 nematodes in the S medium. Before this step, the bacterial culture was washed from the LB broth and resuspended in the S medium until the bacterial concentration reached 0.5 mg/ml. The *E. coli* OP50 cultivation medium consisted of 1000 ml ddH_2_O, 5 g NaCl, 10 g tryptone, and 5 g yeast extract. We used a 96-well plate (Eppendorf, USA) to fill each one with 120 μl of the *C. elegans* + *E. coli* suspension. The plate was sealed with a film and incubated for 48 h in a climatic chamber (Binder, Germany) at 20 °C. To prevent the nematodes from breeding, we added 15 μl of 1.2 mM 5-fluoro-2-deoxyuredin (FUDR) in each well after 48 h and stored at 20 °C for 24 h until the worms reached stage L4. Finally, 15 μl of the tested chemical compounds were added to each well.

To obtain mangiferin, we prepared stocks in 10 mmol/l or 50 mmol/l dimethyl sulfoxide (DMSO). The stock solution was diluted with distilled sterile water to the concentrations of 200 μm, 100 μm, 50 μm, and 10 μm.

We used the scheme below to study the effect of mangiferin on the fat accumulation by *C. elegans* N2 Bristol.

On incubation day 10, the nematodes were subjected to staining [16]. We transferred the samples from the 96-well flat-bottomed plate to a 96-well V-shaped plate and centrifuged at 1000 rpm for 3 min. After that, we removed the supernatant without affecting the nematode sediment and poured 150 μl PBS into each well. The washing stage also included a double three-minute centrifugation at 1000 rpm. The obtained precipitate was mixed with 150 μl of 40% isopropanol as a fixing solution, and the plate underwent 20 min of incubation at room temperature and 3 min of centrifugation at 1000 rpm. Next, we added 150 μl of a borondipyrromethene fluorophore (BDP) 505/515 solution (Lumiprobe, Russia) into each cell to stain the fat (2 μl/ml of stock solution in 40% isopropanol). The incubation, accompanied by gentle stirring, lasted 15 min at room temperature. The plates were kept in a dark place because light is known to destabilize the borondipyrromethene fluorophore 505/515. After another three-minute centrifugation at 1000 rpm, the supernatant was removed without affecting the nematode sediment. A triplicate washing involved an M9T solution. To obtain the solution, we added 5 μl Triton X-10050 ml to the M9 medium, which included 1000 ml ddH_2_O, 3 g KH_2_PO_4_, 5 g NaCl, 6 g Na_2_HPO_4_, and 1 ml of 1 M MgSO_4_. After the washing, the nematodes were mixed with 100 μl M9T, and the stained lipids were scanned using an ImageXpress Mico XL automated fluorescence microscope (Molecular Devices, USA).

The obtained images made it possible to perform a quantitative analysis of body fat accumulated by *C*.*elegans* stained with BDP 505/515 fluorescent dye. After processing, the digital images of stained nematodes were normalized according to the fluorescence intensity.

### 1) Obtaining the images of nematodes stained with BDP 505/515 lipid stain

The experiment involved an ImageXpress Mico XL high-performance automated microscope (Molecular Vevices, USA), which is often used for multiparametric phenotypic screening and analysis of digital images of 96-well plates. The microscope works has a special MetaXpress software that controls the procedure and processes the results. The parameters of the high-speed imaging included: a dry lens with a 4x magnification in transmitted visible light; Cy3 fluorescent dye with 548 nm of maximal absorption and the maximal emission at 562 nm; extinction coefficient of 150,000 1/(mol cm). This equipment made it possible to register the quantitative changes in the level of fat in nematodes on mangiferin incubation day 10.

### 2) Image processing

The image analysis relied on CellProfiler 4.2.1 [17]. The pre-processing included normalizing uneven illumination and removing artifacts, e.g., well edges. Next, the nematodes and their clusters were separated from the background by segmentation in each local area of the sample. Such processing provided highly accurate images of individual objects, both in the fluorescent mode and in the phase contrast mode, regardless of the difference in color intensity.

### 3) Stain normalization

The digital data were represented as an array of values for the glow intensity of the stained fat, where each intensity value was a pixel. Color depth was expressed in bits or pixels: the larger the bits/pixel, the greater the color depth. The spots of stained fat provided the information about the true signal and the background values. The true signal value was obtained by subtracting the background mean. The true signal is the fluorescence intensity caused by the staining of fat on the Cy3 channel; the background signal is the fluorescence intensity associated with the sticky surface of any intracellular debris.

### 4) Normalizing the fluorescence intensity

The purpose of normalization was to correct the differences in the total fluorescence intensity of the entire nematode population after sextuplication. The problem is that the number of nematodes could differ significantly from the number of control animals. Non-normalized data might distort the true value of Cy5. The normalized numerical values of the nematode images made it possible to analyze the inhibitory activity of mangiferin against fat accumulation in nematodes.

## Results and discussion

The initial stage of the experiment included counting the fixed nematodes incubated in mangiferin since not all nematodes were selected to quantify fat accumulation. Figure 2 shows the population of the fixed nematodes accepted for further processing.

**Figure 2.**
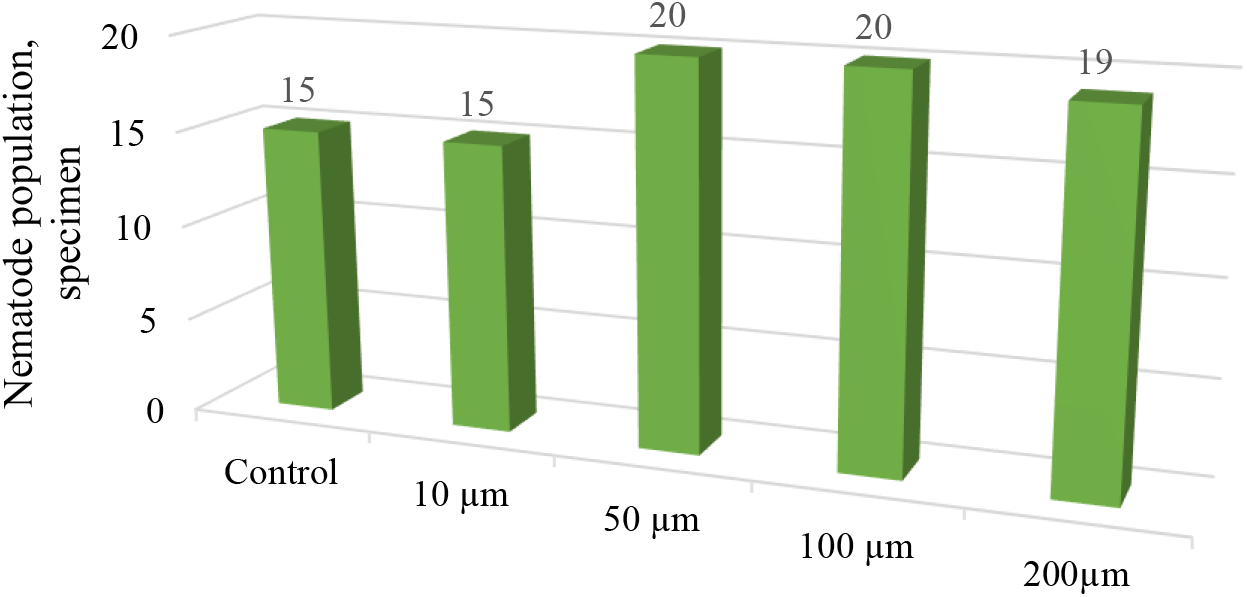
Population of fixed nematodes incubated in mangiferin and accepted for further processing

Apparently, 50-200 μm mangiferin increased the number of nematodes.

CellProfiler 4.2.1 converted the nematode images into digital values of the median fat fluorescence intensity.

The nematode population differed under various test conditions. Thus, the values of the median intensity of fat fluorescence per nematode had to be normalized at the final stage of the research. The data were represented as a histogram of the fat median intensity for each test vs. control. Figure 3 illustrates the effect of mangiferin on fat accumulation in *C. elegans* N2 Bristol.

**Figure 3.**
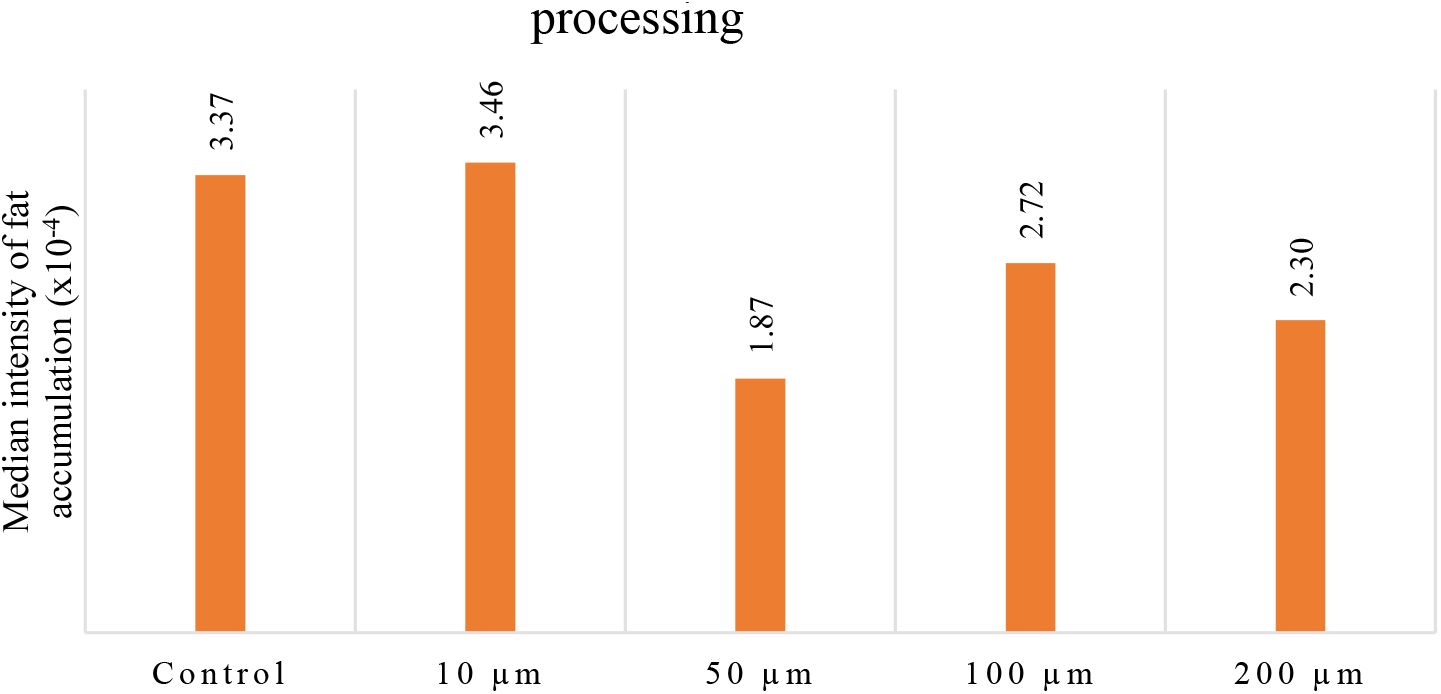
Effect of mangiferin from *Hedysarum neglectum* L. root culture extract on fat accumulation in *C*.*elegans* N2 Bristol

Thus, mangiferin obtained from *Hedysarum neglectum* L. inhibited fat accumulation in nematodes. At 50 μm mangiferin, the decrease in fluorescence was 44.5%; at 100 μm, it was 31.8%; at 200 μm, it was as low as 19.7% (Figure 3).

## Conclusion

This *in vivo* experiment featured the effect of mangiferin isolated from *Hedysarum neglectum* L. root culture on fat accumulation in *Caenorhabditis elegans* N2 Bristol. Mangiferin showed anti-fat activity at a concentration of 50-200 μm. The maximal fat reduction occurred at 50 μm mangiferin, which proves that mangiferin can inhibit fat accumulation in living organisms.

## Funding

The research was part of state assignment on Plant Polyphenols of the Siberian Federal District: Molecular and Spatial Structure, Biofunctional Properties, and Toxicological Safety Indicators in Model Systems *In Vivo*, Project FZSR-2023-0002.

The experiments were conducted on the premises of the Center for Collective Use: Instrumental Methods of Analysis in Applied Biotechnology, Kemerovo State University.

## Notes

### Competing Interest Statement

The authors have declared no competing interest.

## References

1. Moskalev A, Guvatova Z, Lopes IA, Beckett CW, Kennedy BK, De Magalhaes JP, Makarov AA. Targeting aging mechanisms: pharmacological perspectives. Trends Endocrinol Metab. 2022 Apr;Apr(4):266–280. doi: 10.1016/j.tem.2022.01.007.

2. Galikova M, Klepsatel P. Obesity and Aging in the Drosophila Model. Int J Mol Sci. 2018 Jun 27;19(7):1896. doi: 10.3390/ijms19071896

3. Santos AL, Sinha S. Obesity and aging: Molecular mechanisms and therapeutic approaches. Ageing Res Rev. 2021 May;67:101268. doi: 10.1016/j.arr.2021.101268.

4. Jones KT, Ashrafi K. Caenorhabditis elegans as an emerging model for studying the basic biology of obesity. Dis Model Mech. 2009 May-Jun;2(5-6):224–9. doi: 10.1242/dmm.001933.,

5. Relevance of Bioassay of Biologically Active Substances (BAS) with Geroprotective Properties in the Model of the Nematode Caenorhabditis Elegans in In Vivo Experiments / L. S. Dyshlyuk, A. I. Dmitrieva, A. Yu. Prosekov [et al.] // Current aging science. – 2022. – Vol. 15, No. 2. – P. 121–134. –DOI 10.2174/1874609814666211202144911.

6. Park HH, Jung Y, Lee SV. Survival assays using Caenorhabditis elegans. Mol Cells. 2017 Feb;Feb(2):90–99. doi: 10.14348/molcells.2017.0017.

7. Yue Y, Li S, Shen P, Park Y. Caenorhabditis elegans as a model for obesity research. Curr Res Food Sci. 2021 Oct 1;4:692–697. doi: 10.1016/j.crfs.2021.09.008.

8. Queen BL, Tollefsbol TO. Polyphenols and aging. Curr Aging Sci. 2010 Feb;Feb(1):34–42. doi: 10.2174/1874609811003010034.

9. Aloo SO, Ofosu FK, Kim NH, Kilonzi SM, Oh DH. Insights on Dietary Polyphenols as Agents against Metabolic Disorders: Obesity as a Target Disease. Antioxidants (Basel). 2023 Feb 8;12(2):416. doi: 10.3390/antiox12020416.

10. Mayneris-Perxachs J., Alcaide-Hidalgo J.M., de la Hera E., del Bas J.M., Arola L., Caimari A. Supplementation with biscuits enriched with hesperidin and naringenin is associated with an improvement of the Metabolic Syndrome induced by a cafeteria diet in rats. J. Funct. Foods. 2019;61:103504. doi: 10.1016/j.jff.2019.103504

11. Antimicrobial and antioxidant activity of Panax ginseng and Hedysarum neglectum root crop extracts / L.S. Dyshlyuk, N.V. Fotina, I.S. Milentyeva et al // Brazilian Journal of Biology. – 2022. –V. 84. https://doi.org/10.1590/1519-6984.256944

12. Sekiguchi Y, Mano H, Nakatani S, Shimizu J, Kataoka A, Ogura K, Kimira Y, Ebata M, Wada M. Mangiferin positively regulates osteoblast differentiation and suppresses osteoclast differentiation. Mol Med Rep. 2017 Aug;Aug(2):1328–1332. doi: 10.3892/mmr.2017.6752.

13. Imran M, Arshad MS, Butt MS, Kwon JH, Arshad MU, Sultan MT. Mangiferin: a natural miracle bioactive compound against lifestyle related disorders. Lipids Health Dis. 2017 May 2;16(1):84. doi: 10.1186/s12944-017-0449-y.

14. Rodríguez J, Di Pierro D, Gioia M, Monaco S, Delgado R, of Coletta M, Marini S. Effects a natural extract from Mangifera indica L, and its active compound, mangiferin, on energy state and lipid peroxidation of red blood cells. Biochim Biophys Acta. 2006;1760(9):1333–42.

15. Effects of bioactive substances isolated from Siberian medicinal plants on the lifespan of Caenorhabditis elegans / E. R. Faskhutdinova, A. S. Sukhikh, V. M. Le [et al.] // Foods and Raw Materials. – 2022. – Vol. 10, No. 2. – P. 340–352. –DOI 10.21603/2308-4057-2022-2-544.

16. Kuzmic M, Javot H, Bonzom J-M, Lecomte-Pradines C, Radman M, Garnier-Laplace J, Frelon S. In situ visualization of carbonylation and its co-localization with proteins, lipids DNA and RNA in Caenorhabditis elegans // Free Radical Biology and Medicine. – 2016. –№11.–p.465–474.https://doi.org/10.1016/j.freeradbiomed.2016.11.004

17. Label-free imaging of lipids depositions in C. elegans using third-harmonic generation microscopy / G.J. Tseverelakis, E.V. Megalou, G. Filippidis et al. // PLOS One. – 2014. – Vol. 9. – P. e84431. https://doi.org/10.1371/journal.pone.0084431

